# Interactions between tick-borne encephalitis virus non-structural protein 1 and blood-brain barrier tight junction proteins: potential clues to strain-specific neuropathogenicity

**DOI:** 10.1101/2025.08.12.669603

**Authors:** Jason Pun, Jarinya Chaopreecha, Jordan Liburd, Shahid Khalid, Ian M Jones, Jennifer H. Tomlinson, Niluka Goonawardane

## Abstract

Tick-borne encephalitis virus (TBEV) invades the central nervous system (CNS) through strain-specific mechanisms that remain poorly understood. In mosquito-borne orthoflaviviruses such as dengue and yellow fever viruses, the non-structural protein 1 (NS1) has been shown to disrupt endothelial barrier integrity by targeting tight junction proteins (TJPs), thereby facilitating viral neuroinvasion. However, comparable mechanisms in TBEV remain largely unexplored.

Here, we investigate the potential interaction of NS1 from high (Hypr)- and low (Vs)-pathogenicity TBEV strains to different blood-brain barrier (BBB) TJPs, using AlphaFold3 (AF3) multimer modelling and *in vitro* binding assays. We find that NS1 from the highly pathogenic strain exhibits higher predicted interactions with multiple TJPs, including two junctional adhesion molecules (JAM-A, JAM-B) and Claudin-10, which are critical component of the paracellular barrier. In contrast, low pathogenic strain Vs interaction was limited to JAM-A and Claudin-5.

Experimental validation using recombinant NS1 proteins revealed strain-specific binding profiles: Hypr NS1 displayed high-affinity, saturable direct binding to immobilized JAM-A (KD = 0.271 µg/mL), whereas Vs NS1 showed negligible interaction (KD = 0.000023 µg/mL). No binding to ZO-1, a barrier scaffold lacking an extracellular domain, was observed for either strain. This differential interaction may be modulated by 22 specific amino acid substitutions localized to the Wing and β-ladder domains, which distinguish the highly neurovirulent Hypr strain from the avirulent Vs strain. Notably, despite its weak JAM-A interaction, the Vs strain is associated with slow-progressing infections that can culminate in chronic neurological disease, highlighting the need for further investigation into noncanonical pathways of neuroinvasion.

**Graphics:** *A graphical illustration of low-and high-pathogenic TBEV mediated compromised blood-brain barrier (BBB):* The left panel illustrates the cellular architecture of the BBB, including pericytes, endothelial cells, astrocytes, and blood vessels. The central panel depicts an intact BBB with tight junction proteins (JAM-A, ZO-1, Claudin) maintaining barrier integrity. The right panel shows strain-specific pathogenesis, highlighting a direct interaction between NS1 from the highly pathogenic Hypr strain and JAM-A, which may contribute to differential neuroinvasive outcomes. The schematic was created using BioRender.com. 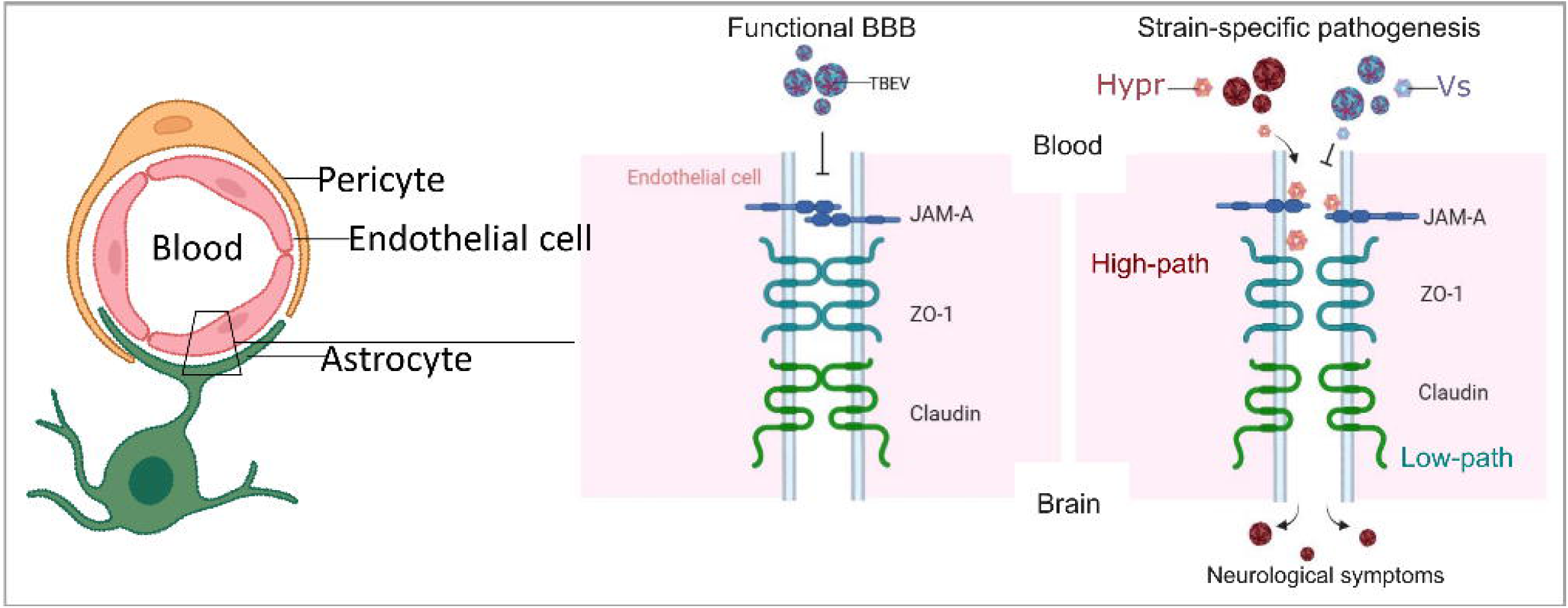

## Introduction

The NS1 is a structurally conserved critical virulence factor encoded by orthoflaviviruses, including TBEV. NS1 is a glycoprotein that exists in multiple forms – intracellular, membrane-associated, and secreted (Figure. 1A) – all of which contribute to the virus’s replication, immune evasion, and pathogenesis ^1–3^. While the role of NS1 has been extensively studied in mosquito-borne orthoflaviviruses such as dengue virus (DENV) ^4, 5^, yellow fever virus (YFV) ^6, 7^ and Zika virus (ZIKV) ^8, 9^, the functions and host interactions of TBEV NS1 remain less well characterised, particularly in the context of strain-specific clinical outcomes.

**Figure 1.**
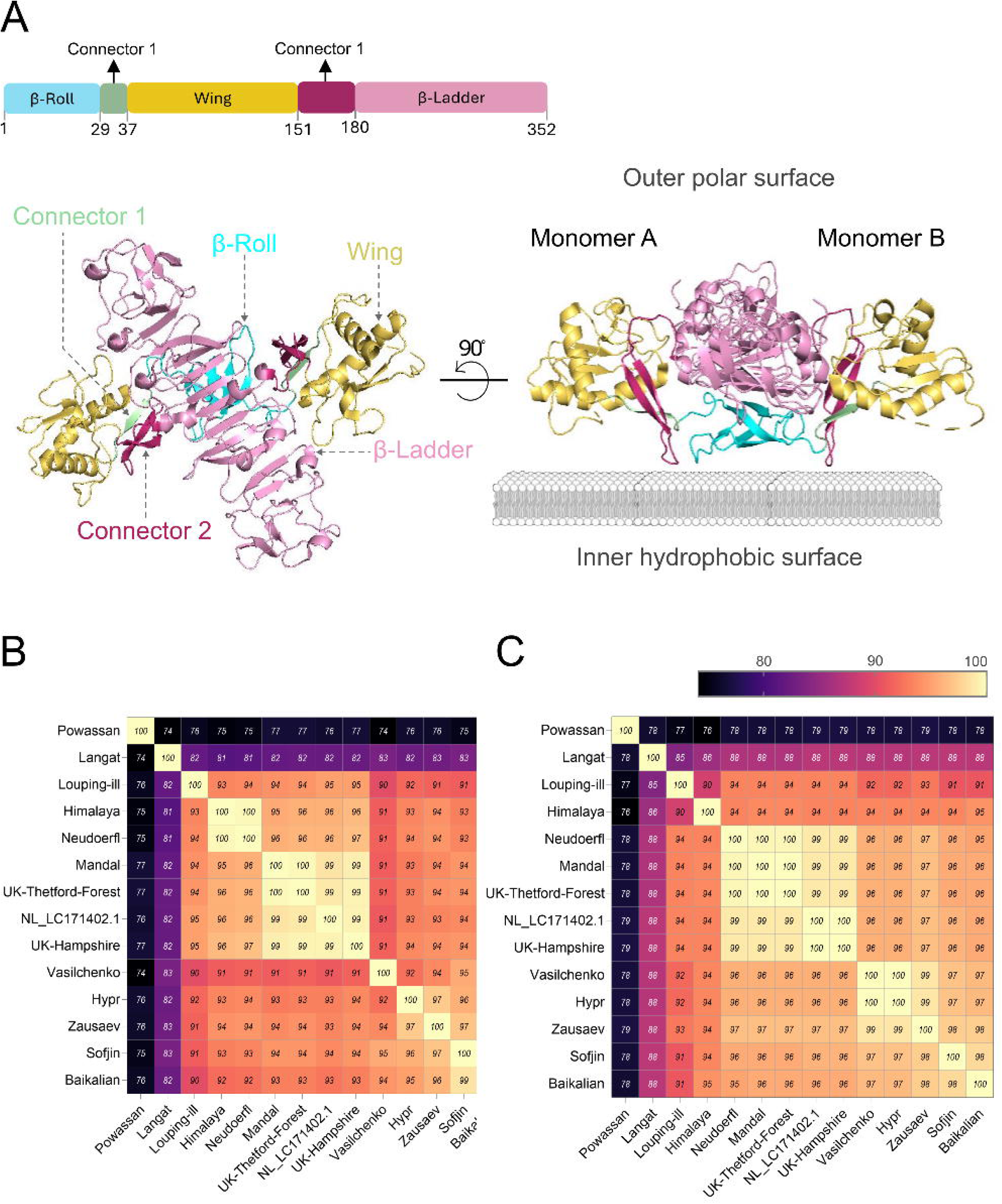
Cryo-EM structure of orthoflavivirus non-structural protein 1 (NS1). **(A)** The NS1 dimer is shown in cartoon representation. The ribbon diagram highlights the β-roll, wing, β-ladder, and connector domains (1 and 2), which are coloured blue, yellow, pink, green, and purple, respectively, in one monomer. The dengue virus (DENV) NS1 dimer (PDB ID: 4OIG) is shown interacting with a phospholipid membrane. **(B)** Percent identity matrix for the NS1 **(B)** or **(C)** (envelop) E proteins among the indicated 14 tick-borne Orthoflaviviruses (TBFV), generated by ClustalW. The each in the matrix represents the percentage of identical residues between each pair of sequences in the alignment, calculated by the aligned positions. The identity values are shown as an inferno heatmap with the gradient from highest (light yellow; 100%) to lowest (dark purple; <74%).

**Figure 2.**
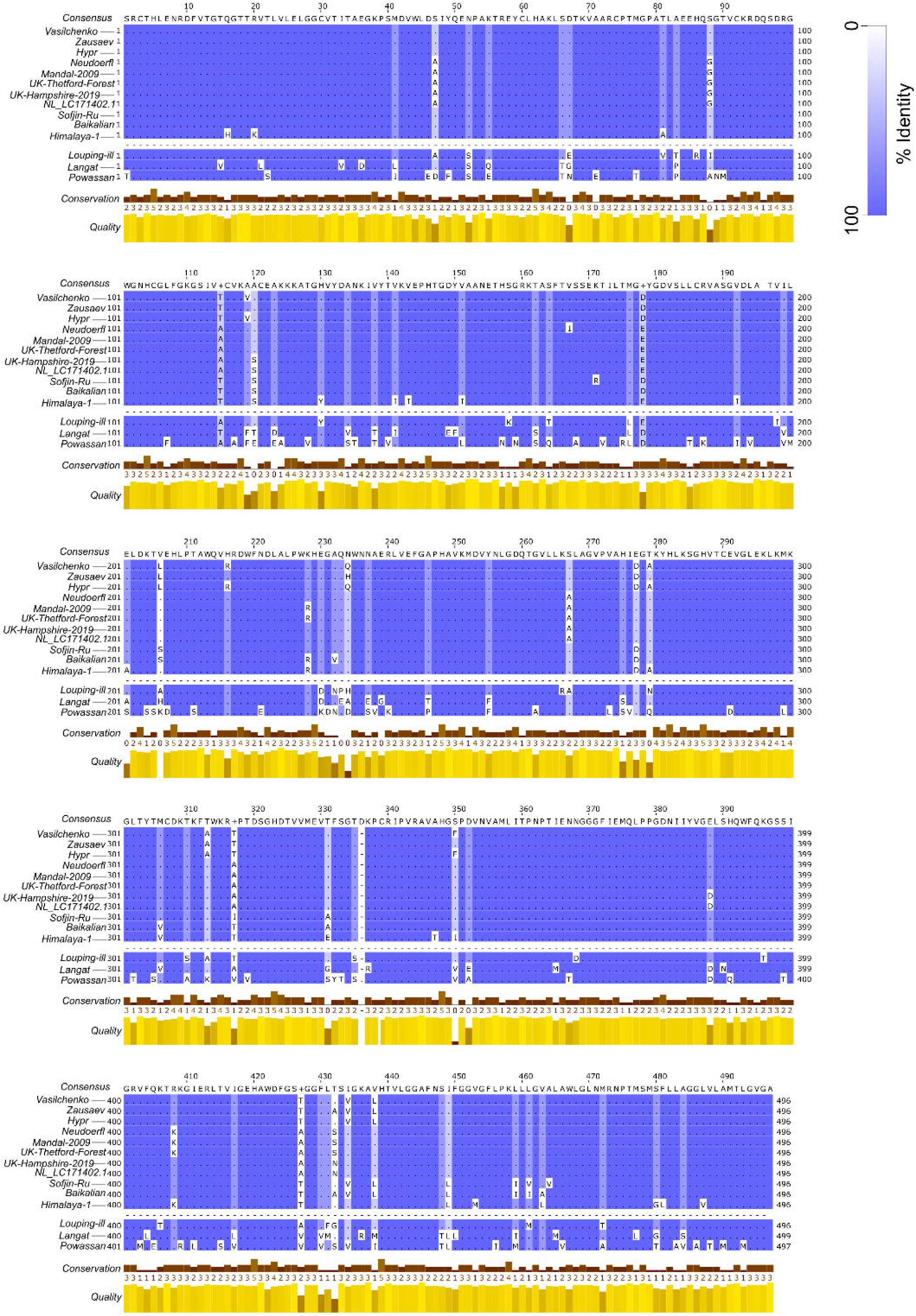
Amino acid sequence alignment of the NS1 protein from tick-borne Orthoflaviviruses (TBFV). The NS1 proteins of different TBFV were compared using a multiple alignment tool, CLUSTAL Omega. The analysis showed conserved regions among the NS1 sequences of these viruses and indicated as percentage (%) sequence identity form highlighted areas of similarity (100%; dark purple) to dissimilarity (0; white). The consensus for all 14 sequences is indicated at the top. The alignments are grouped into TBEV isolates (Vasilchenko [accession number; AF069066.1], Hypr [U39292.1], Neudoerfl [U27495.1], Zausaev [AF527415.1], UK-Thetford-Forest [MN128700.1], UK-Hampshire-2019 [MN661145.1], Sofjin-Ru [JN229223.1], Mandal-2009 [KF991107.1], NL [LC171402], Baikalian [EF469661.1], Himalaya-1[MG599476.1] and other closely related TBFV (Louping-ill-virus [Y07863.1], Langat-virus_[NC_003690.1] and Powassan [NC_003687.1]. The sequence conservation of total alignments less than 20% of gaps and BLOSUM62 alignment quality is shown.

TBEV is a neurotropic arbovirus that can cause severe neurological disorders ^10, 11^, including encephalitis and meningitis ^12–14^, often leading to long-term neurological sequelae ^15, 16^. A defining feature of TBEV pathogenesis is its capacity to breach the BBB and trigger neuroinflammation in a strain-dependent manner ^17, 18^. Recent evidence implicates the NS1 as a key virulence factor in endothelial dysfunction and immune modulation ^4–7^. In related viruses (including dengue and yellow fever virus), secreted NS1 (sNS1) has been shown to compromise vascular integrity by targeting tight junction components such as ZO-1 and several claudins, thereby promoting vascular leakage and amplifying inflammatory responses^19^. Despite these parallels, the contribution of strain-specific TBEV NS1 to BBB disruption remains poorly defined, and direct mechanistic insights are lacking. Addressing these gaps is critical to unravelling the pathogenesis of TBEV and developing targeted therapies.

In this study, we investigated the potential interactions between the NS1 protein of high- and low-virulence TBEV strains and a series of TJPs to explain the observed functional consequences at the BBB. Using AlphaFold3 (AF3) multimer model ^20, 21^, an advanced deep learning framework that significantly enhances the accuracy of multimeric protein complex prediction, we identified and validated strain-specific binding patterns of NS1 to TJPs. These findings provide mechanistic insights into how NS1 may contribute to BBB disruption during the neurological phase of TBEV infection and highlight a potential therapeutic avenue to mitigate neuroinvasion and subsequent neuropathology by targeting this interaction axis.

## Method

### Recombinant TBEV NS protein generation and purification

N-terminally His-tagged Vs (accession number: AF069066.1) or Hypr (accession number: U39292.1) NS1 was heterologously expressed in *E. coli* BL21 Gold cells harbouring the pET-28a-TEV-NS1 plasmid (synthesised by Genscript). Protein was expressed in LB medium containing 50 μg/ml kanamycin. Briefly, cultures were incubated at 37 °C, with shaking at 200 rpm until an OD_600nm_ of ∼0.6 was reached. Protein expression was induced with 1 mM IPTG and cultures were incubated at 25 °C overnight with shaking at 200 rpm. Cells were then harvested by centrifugation at 4500 xg for 30 min at 4 °C. Cell pellet was subsequently resuspended in 50 mM TrisHCl, 500 mM NaCl, 2 mM MgCl_2_, 10 mM imidazole, 10 % v/v glycerol pH 8.0 containing benzonase (Merck) and cOmplete EDTA-free protease inhibitors (Roche) and lysed by sonication on ice before clearing lysate by centrifugation at 11,000 xg at 4 °C for 20 min. NS1 however was located in the insoluble fraction and was recovered via on-column refolding as follows. The insoluble fraction was resuspended in denaturing buffer (20 mM TrisHCl, 300 mM NaCl, 20 mM imidazole, 5 M guanidine HCl, 10 mM DTT, pH 8.0) and incubated on ice for 1 hour before centrifugation at 13,000 xg for 20 min at 4 °C. Extracted protein was then applied to a Ni-NTA column (Qiagen) and washed with denaturing buffer. A gradient wash was then applied, gradually reducing the guanidine HCl and DTT concentrations to 0 M and the column was further washed in refolding buffer (20 mM TrisHCl, 300 mM NaCl, 20 mM imidazole, pH 8.0). Protein was eluted in elution buffer (20 mM TrisHCl, 300 mM NaCl, 500 mM imidazole, pH 8.0) and purity was assessed using SDS-PAGE. Purified NS1 protein was exchanged into 50 mM TrisHCl, 500 mM NaCl, 2 mM MgCl_2_, 10 mM imidazole, 10 % v/v glycerol pH 8.0 and stored at −80 °C. Successful refolding was assessed using CD spectroscopy, comparing with commercially available NS1.

### Multiple sequence alignment and percent identity matrix generation

NS1 protein sequences of TBEV isolates and closely related tick-borne Orthoflaviviruses were downloaded from the NCBI (Vasilchenko [accession number; AF069066.1], Hypr [U39292.1], Neudoerfl [U27495.1], Zausaev [AF527415.1], UK-Thetford-Forest [MN128700.1], UK-Hampshire-2019 [MN661145.1], Sofjin-Ru [JN229223.1], Mandal-2009 [KF991107.1], NL [LC171402], Baikalian [EF469661.1], Himalaya-1[MG599476.1] and other closely related TBFV (Louping-ill-virus [Y07863.1], Langat-virus_[NC_003690.1] and Powassan [NC_003687.1]) and aligned using ClustalW, implemented in BioEdit. Default parameters were used unless otherwise specified, including the Gonnet substitution matrix, a gap opening penalty of 10, and a gap extension penalty of 0.2. Following alignment, a Percent Identity Matrix (PIM) was generated, which calculates the pairwise percentage of identical residues between aligned sequences. This matrix provides a quantitative measure of sequence similarity and was used to assess evolutionary relationships and sequence conservation.

### NS1-BBB TJP interaction prediction and interface analysis by AlphaFold3 (AF3)

The predicted interaction between TBEV Hypr or Vs NS1 and the host TJPs was modelled using AF3, following the approach described by Su et al., 2024 ^22^. The AF3 structural prediction was visualised using PyMOL ^23^ (https://pymol.org/2/). Briefly, to identify interface residues, the NS1–TJP complex was loaded into PyMOL, and residues at the protein–protein interface were selected manually based on proximity and surface contact. The “find polar contacts” function was used to detect potential hydrogen bonds and salt bridges between the two proteins. Interacting residue pairs were further refined by selecting only those forming direct contacts between NS1 and TJP.

Distances between interacting atoms were measured using the “distance” measurement tool in PyMOL. All interacting residue pairs were ranked based on the shortest interatomic distances. The top-ranking interactions–those with the shortest and most structurally significant contacts were visualized and annotated in the PyMOL structural model. A complete list of all detected interactions, including residue identities and distances, is provided in Tables 4-6.

### Molecular docking and thermodynamic analysis

Ns1–TJP docking was performed using the HDOCK web server (http://hdock.phys.hust.edu.cn/), which uses a hybrid scoring function combining template-based modelling and free docking algorithms. Homodimers of NS1 proteins from TBEV Hypr and Vs strains were docked against several human full-length or their extracellular domains, including Claudin-5, Claudin-10, JAM-A, JAM-B, and JAM-C. Docking scores were recorded as a proxy for binding energy, with higher negative scores indicating stronger predicted interaction. The docking was performed in default mode, allowing the server to automatically identify potential binding sites and generate a ranked list of predicted complex models.

To estimate binding affinity under physiological (25°C, 298.15 K) and febrile (40°C, 313.15 K) conditions, Gibbs free energy (ΔG) and equilibrium dissociation constants (Kd) were calculated from docking scores using the thermodynamic equation:

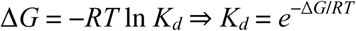

where R is the gas constant (0.001987 kcal·mol¹·K¹), and T is temperature in Kelvin. We assumed docking scores approximate ΔG in kcal/mol, consistent with prior benchmarks for HDOCK scoring. Fold changes in Kd between 25°C and 40°C were computed to assess temperature sensitivity (Table 2).

### Enzyme linked immunosorbent assay (ELISA) for NS1–JAM-A/ZO-1 interaction

High-binding 96-well plates were coated with recombinant human JAM-A (Kactus, HM10A) or ZO-1 (Invitrogen, RP-99371) at a concentration of 1 µg/mL in 0.1 M sodium bicarbonate buffer (pH 9.6) and incubated for 2 h at room temperature (RT). Wells were washed three times with PBS containing 0.1% Tween-20 (PBS-T), followed by blocking with 5% bovine serum albumin (BSA) in PBS supplemented with 0.1 mM EDTA for 1 hour at RT. Recombinant NS1 proteins from TBEV Hypr or Vs strains were serially diluted 2-fold and incubated for 2 h at RT. After washing (3× with PBS-T), bound NS1 was detected using a mouse anti-TBEV NS1 monoclonal antibody (R&D Systems, MAB10106; 0.05 µg/mL), followed by incubation with HRP-conjugated anti-mouse IgG (Sigma, A9044; 1:40,000) for 1[h at RT. Plates were washed again and incubated with TMB peroxidase substrate (Sigma, T5569) in the dark for up to 30[minutes at RT. The reaction was stopped with 1[M hydrochloric acid, and absorbance was measured at 450 nm using a NanoQuant Infinite M200 plate reader (Tecan).

For ELISA data processing and curve fitting, raw OD 450nm values were averaged across technical replicates and plotted against the corresponding NS1 concentrations. Binding curves were fitted using a one-site binding hyperbola model:

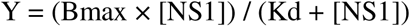

where Y is the OD450 signal, Bmax is the maximum binding, and Kd is the dissociation constant. Nonlinear regression was performed using the curve-fit function from the SciPy library in Python. The fitted parameters (Bmax and Kd) were extracted for each strain and experiment.

For statistical analysis, selected concentrations (0.01 µg/mL and 0.25 µg/mL), OD450 values were compared between Hypr and Vs NS1 using unpaired two-tailed Student’s t-tests. P-values were calculated using the ‘ttest_ind’ function from SciPy. Statistical significance was annotated on bar plots.

All plots were generated using Matplotlib in Python. Binding curves were displayed with raw data points (mean ± SD) and fitted curves overlaid. Bar graphs were used to visualise OD450 nm values at selected concentrations, with error bars representing standard deviation.

## Results

### Relative diversity of NS1 protein among tick-borne orthoflaviviruses

To investigate the evolutionary relationships among tick-borne orthoflaviviruses (TBFVs), we generated pairwise percentage identity matrices based on NS1 protein sequences from 14 viral Isolates. This panel included multiple TBEV strains, including recent isolates from the UK (Thetford-Forest [MN128700.1], Hampshire-2019 [MN661145.1]) and the Netherlands (NL [LC171402]), as well as more distantly related viruses, including Powassan virus (POWV; NC_003687.1), Langat virus (LGTV; NC_003690.1), and Louping-ill virus (LIV; Y07863.1). The resulting identity values, visualised as a heatmap (Figure 1B), revealed high conservation within the broader TBEV group, with pairwise identities generally exceeding 92%. In contrast, interspecies comparisons showed substantially lower identity values (e.g., 77–83% between POWV and TBEV strains), consistent with their divergent evolutionary trajectorie.

In contrast, the envelope (E) protein, which mediates host cell entry and serves as a major antigenic determinant, exhibited less pronounced subtype-specific divergence (Figure 1C). Within the TBEV group, E protein identities ranged from 94% to 100%, with Far Eastern (Sofjin) subtypes forming a distinct cluster. Notably, Hypr and Vasilchenko strains shared 100% identity, indicating minimal divergence within this subgroup. Comparisons with more distantly related flaviviruses, such as Powassan and Langat virus, revealed lower E protein identities (77–88%), inline with the antigenic profiles. Collectively, these data suggest that while both NS1 and E proteins are conserved within TBEV, NS1 may be subject to distinct selective pressures, potentially reflecting differences in functional constraints or immune evasion strategies.

### Structural comparison of NS1 across flavivirus strains reveals lineage- and pathogenicity-specific amino acid differences

To investigate the structural basis underlying differences in pathogenicity and vector specificity among flaviviruses, we performed a comparative structural analysis of NS1 from representative tick-borne (TBEV) and mosquito-borne (ZIKV, JEV, and WNV) viruses. We focused on identifying lineage-specific amino acid (AA) differences that correlate with pathogenic potential and tick- or misquote-vector tropism.

Similar to other orthoflaviviruses, the NS1 protein of TBEV comprises three distinct domains: a β-roll (residues 1–29), a wing domain (residues 38–180), and a β-ladder domain (residues 181–352) (Figure 1A) ^1, 2^. High-resolution structural alignment of NS1 monomers from the TBEV-Vs (low-pathogenic) and -Hypr (high-pathogenic) strains revealed several discrete (22 AA) substitutions, particularly localised in the surface exposed wing domain (9 AA) and the distal β-ladder (13 AA) domain (Figure. 3A). These domain clustering and variations may enhance host membrane association or modulate innate immune recognition, potentially contributing to the heightened neurovirulence observed in the Hypr strain.

**Figure 3.**
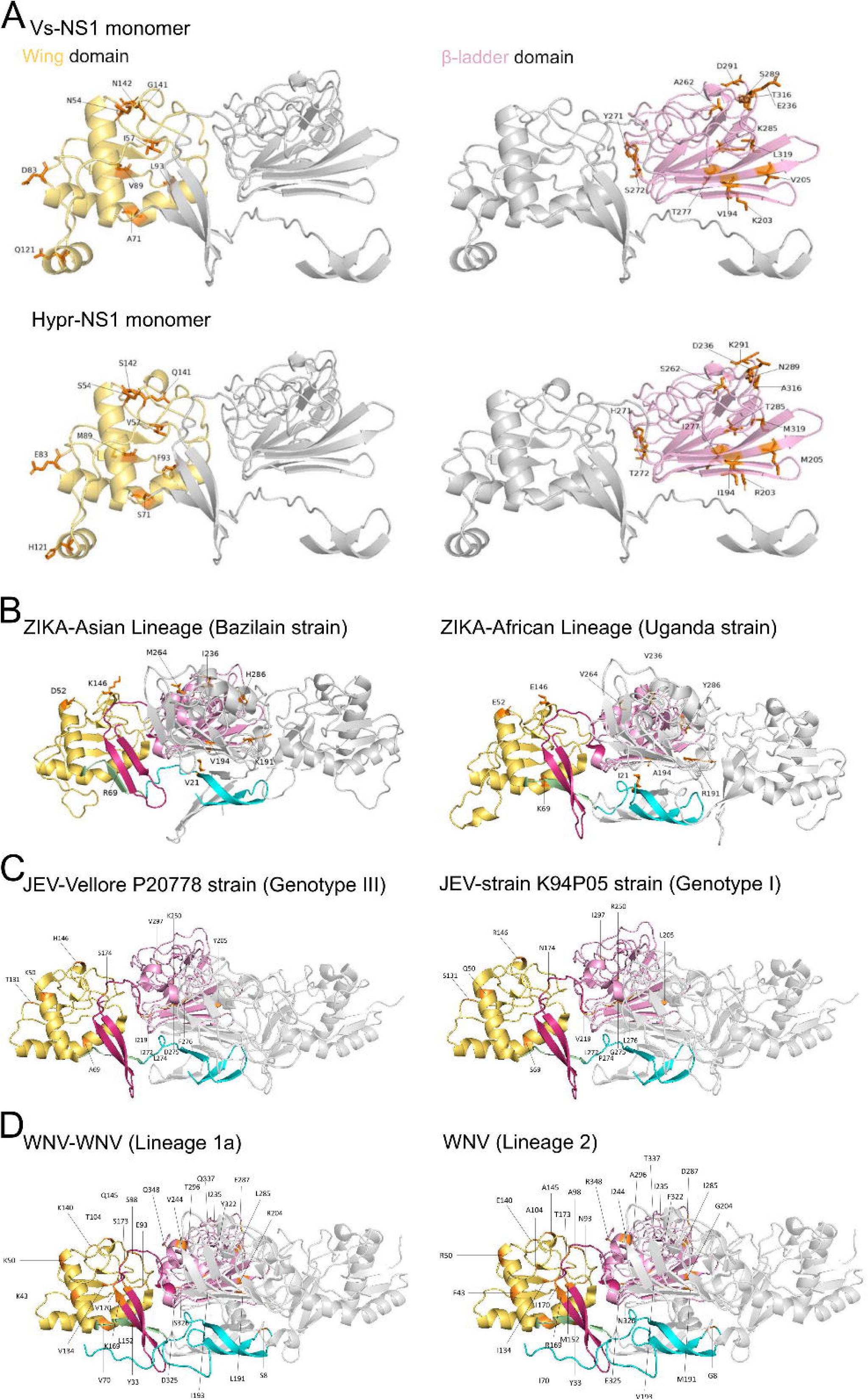
Localisation of amino acid (AA) changes in representative low- and high-pathogenic strains of TBEV (A), ZIKA (B), JEV (C) and WNV (D) NS1. The domains are colour corded as in Figure 1A. Initial atomic positions were based on (PDB ID: 4OIG) for all. **(A)** Position of 22-AA changes in the three-dimensional structure of the Vs (low-path)- and Hypr (high-path)-NS1 monomer prediction by AlphaFold3, visualised in PyMol. Focused view of the mutations in the Wing (yellow) or β-ladder (Pink) domain are indicated. The mutated amino acid is shown in atomic stick representation. **(B-D)** NS1 of high- and low-pathogenic strains are indicated in left and right, respectively. **(B)** Dimer of Asian (PDB: 5GS6) (left) and African (PDB:5K6K)-ZIKA (right) virus NS1, showing all 9 AA changes (orange). The domains are colour corded as in Figure 1A. **(C)** Dimer of JEV genotype III-and genotype I-NS1 with 13 site-specific amino acid changes highlight in orange colour. **(D)** NS1 dimer of lineage 1a- and lineage 2-WNV NS1, indicating 30 site-specific amino acid mutations.

In Mosquito-borne ZIKV virus lineages linked to variable neurovirulence, NS1 structures exhibit lineage-specific features changes in 9 AA. Comparison of NS1 monomers from Asian lineage (Brazil strain) and African lineage (Uganda strain) ZIKV isolates revealed potentially significant AA changes in the central β-roll (1 AA), β-ladder (5 AA) and wing (3 AA) domain (Figure. 3B). While the overall NS1 architecture is conserved, the Asian lineage, which has been epidemiologically linked to congenital Zika syndrome ^24–26^, displayed surface-exposed wing domain substitutions that may impact host protein interactions, NS1 secretion efficiency, or immune modulation. These results suggest that minor structural remodelling in the NS1 surface topology may underlie differences in tissue tropism and pathogenesis.

Structural variation in JEV NS1 from Genotype III (High-pathogenic P20778 strain) and Genotype I (K94P05 strain) exhibited conserved tertiary structures with 13 genotype-specific AA differences in connector 2 loop (1 AA), the wing domain (4 AA) and the β-ladder (8 AA) (Figure. 3C). These residues are positioned in proximity to putative glycosylation and oligomerisation sites (Table 3), potentially influencing NS1 dimerisation or host immune interactions. Given that genotype I strains have displaced genotype III in several endemic regions, these structural distinctions related to the changes favouring transmissibility or immune escape ^27^.

NS1 from WNV lineages shows progressive changes with 30 AA, correlating with emerging pathogenicity. Structural comparison of NS1 from WNV Lineage 1a (highly pathogenic) and Lineage 2 (historically low-pathogenic, now emerging) revealed high conservation overall, with AA substitutions in each domain. The highest number were in the surface exposed (wing) domain (11 AA). 13 AA changes were in the β-roll (Figure. 1D). Notably, Lineage 2 NS1 exhibits substitutions in regions linked to lipid binding and ER membrane association, possibly linked with recent observations of increased neuroinvasive potential in clinical cases^28^. These findings support the hypothesis that NS1 evolution within Lineage 2 may contribute to its changing pathogenic profile. Powassan virus, which is a tick-borne arbovirus endemic to North America, Linage 1 and 2, in comparison, differed in its NS1 sequence in only 1 AA in the β-ladder (Phenylalanine (F) to Leucine (L) at position 247) (Table 3).

These comparative structural analyses demonstrate that lineage- and strain-specific mutations in NS1 cluster in functionally important domains, such as the β-ladder and wing domain, across both tick- and mosquito-borne flaviviruses. These substitutions may influence protein– membrane interactions, immune evasion mechanisms, and tissue tropism, providing a potential molecular basis for differences in virulence and vector specificity. This suggests that adaptive amino acid changes in NS1 contribute to the evolution of arbovirus pathogenicity, particularly in the context of emerging strains with altered clinical outcomes.

### NS1 interacts with tight junction proteins in a strain-specific manner

Given the established role of related arbovirus NS1 in disrupting endothelial integrity, we next investigated whether TBEV NS1 interacts with key TJPs, including JAM-A, JAM-B, JAM-C, claudin-5, and claudin-10. Structural predictions using AlphaFold3-Multimer were performed for both the hypervirulent (Hypr) and avirulent (Vs) NS1 variants, focusing on the extracellular domains (ectodomains) of the TJPs, which represent the biologically accessible interfaces for secreted or membrane-associated NS1. ZO-1, a cytoplasmic scaffold protein lacking an ectodomain, was included as a negative control. Of note, here we applied an iPTM threshold of ≥ 0.6 to define only high-confidence protein-protein interactions, consistent with recent findings by Abulude et al. (2025) that demonstrated that interactions with iPTM scores in this range could be experimentally validated using the BACTH system and corresponded to biologically relevant complexes in *Bdellovibrio bacteriovorus* ^29^

The AF3 models predicted a confident interaction between JAM-A (Figure. 4 A-B) and both Hypr and Vs NS1 (iPTM ≥ 0.6; low interface PAE; Table 1 and Table 4), involving the membrane-distal Ig-like domain of JAM-A engaging a hydrophobic groove on the NS1 dimer.

**Figure 4.**
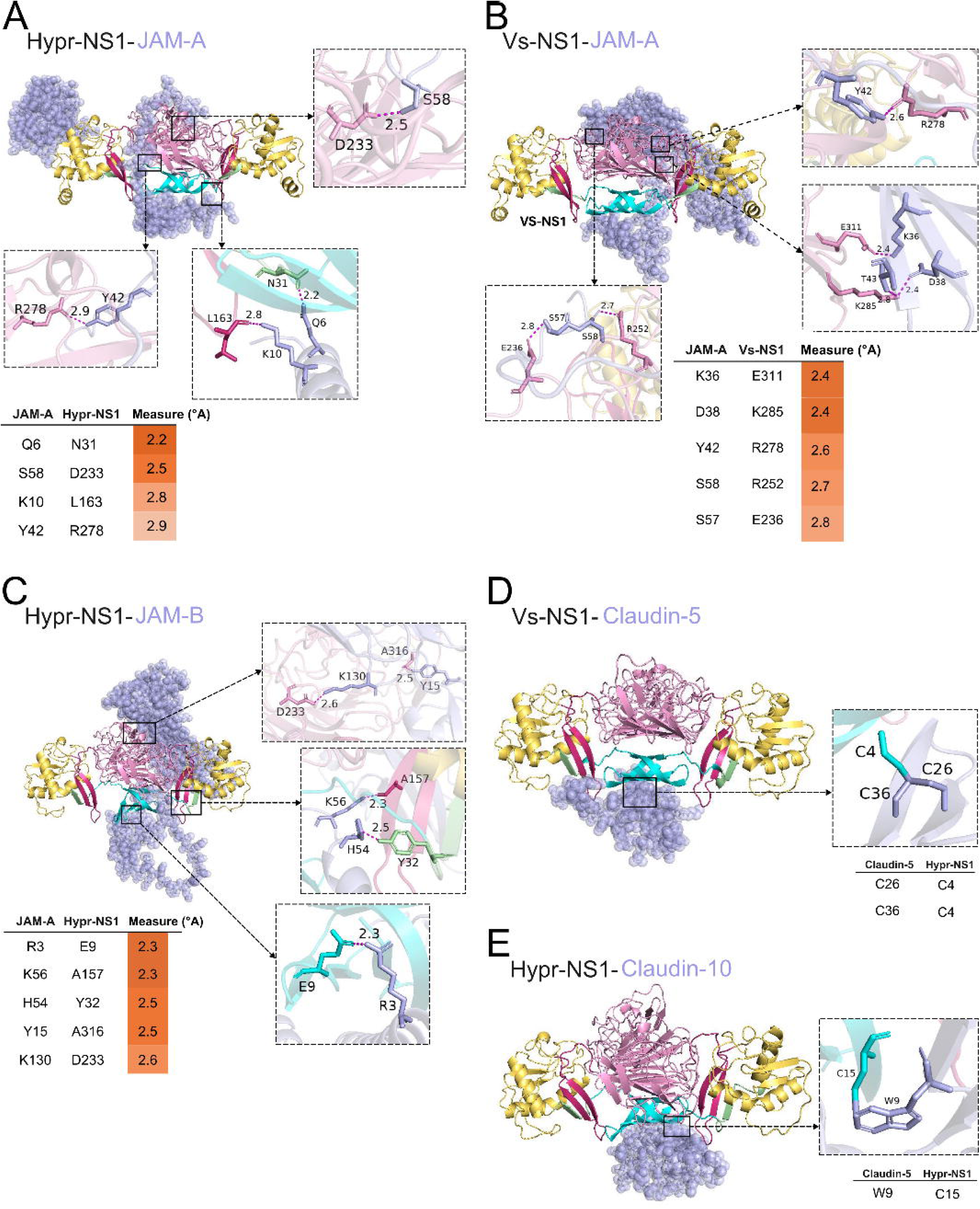
Predicted NS1–TJP interaction models generated by AlphaFold3 (AF3). Interface interacting residues are shown for TBEV Hypr-NS1 **(A)** or Vs-NS1 **(B)** with JAM-A. Hypr-NS1 interactions with JAM-B **(C)** and Vs-NS1 with Claudin 5 **(D)** are indicated. The predicted selected distances (range: ≤2.0–3.3 Å) between amino acid interface residues of NS1 and human TJPs are shown. A heatmap indicates interaction distances, with dark orange representing shorter distances and light orange representing longer ones. NS1 domains are colour coded as in Fig. 1A.

Among other JAM family members, only JAM-B (Figure. 4C, Table 5) showed moderate interaction with Hypr NS1 (iPTM = 0.63), while JAM-C did not interact with either variant, indicating selective engagement rather than a general JAM-binding phenotype. As expected, no direct interactions were predicted between NS1 and ZO-1 (Table 1), consistent with its intracellular localization and supporting the specificity of the modelling approach ^30^.

Claudin-5, which contains two short extracellular loops (ectodomain 1 and 2), similar to Claudin-10, did not form stable complexes with Hypr NS1 (Table 1 and table 6). However, Vs NS1 showed a modest interaction with claudin-5 (Figure 4D), whereas only Hypr-NS1 (Figure 4D) was predicted to form interactions with Claudin-10 (Table 1, ipTM = 0.64, pTM = 0.75), suggesting potential strain-specific differences in claudin engagement. Parallel molecular docking of TBEV NS1 proteins (from Vs and Hypr strains) to claudin-5 and claudin-10 were performed using HDOCK server, modelling both ectodomain 1 and 2 separately.

Based on HDOCK’s reliability thresholds, only interactions with docking scores >200 are firm interactions. Similar to AF3 predictions, this shows the interaction between Vs-NS2 and claudin-5, as well as Hypr-NS1 and claudin-10. Notably, both interactions involved residues within ectodomain 1 (Table 2), suggesting a conserved structural interface that may underpin selective junctional targeting and contribute to modulation of endothelial barrier integrity. These findings support a model in which claudin disruption during flavivirus infection is likely virus-specific and/or indirect ^31^, mediated via signalling downstream of JAM-A or TLR4 activation.

To validate these predictions, we performed ELISA-based binding assays using recombinant NS1 and immobilised JAM-A or ZO-1. Hypr NS1 bound JAM-A in a dose-dependent manner (KD = 0.2714 µg/mL), whereas Vs NS1 showed negligible binding across all tested concentrations (KD ≤ 0.0006 µg/mL) (Figure. 5A–C). At both 0.1 and 0.25 µg/mL, Hypr NS1 exhibited significantly higher optical density (OD450 nm) than Vs NS1 (Figure. 5D), confirming a consistent and measurable difference in binding affinity.

**Figure 5.**
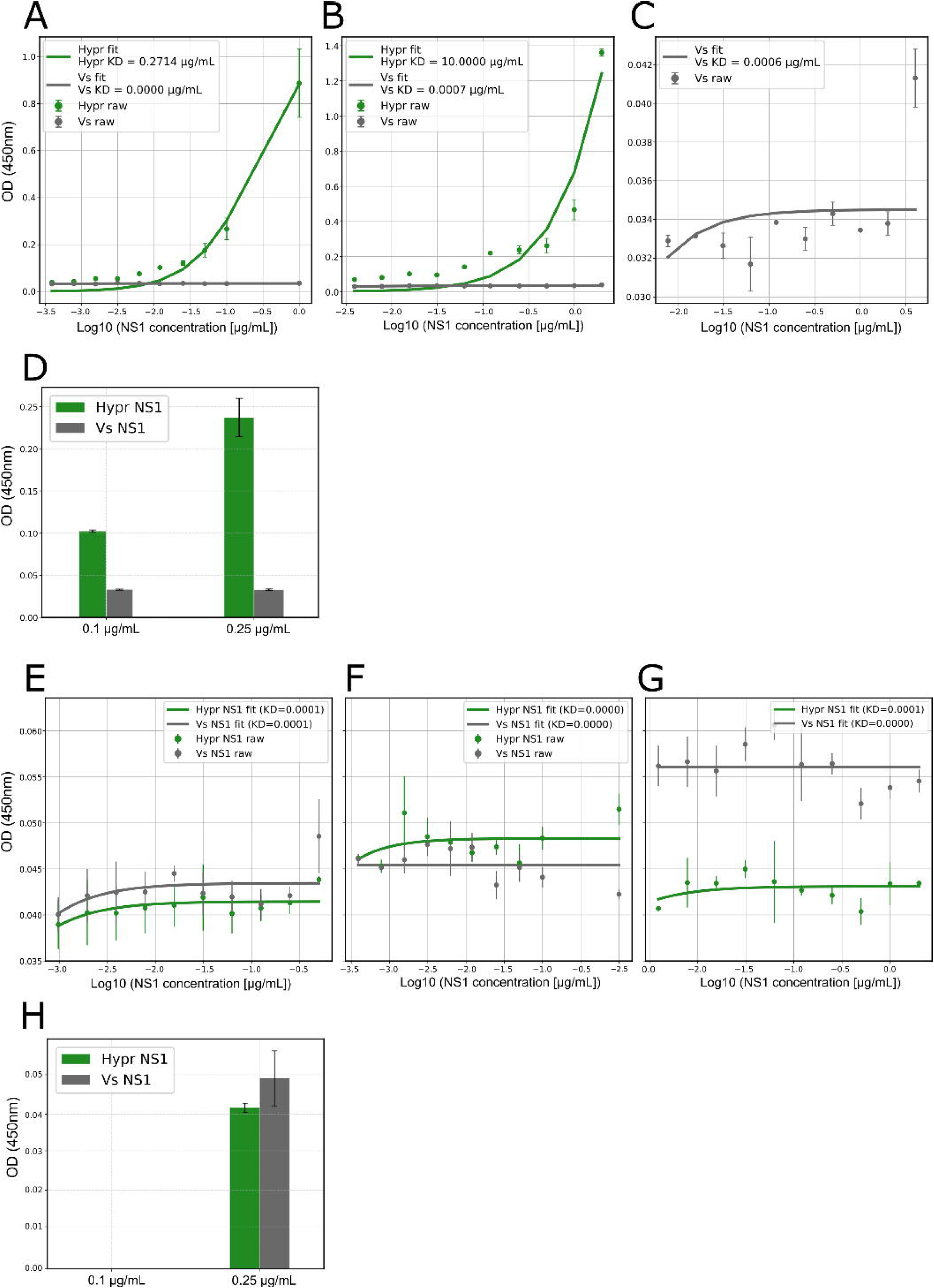
Strain-specific binding of TBEV NS1 to human TJPs. **(A–C)** Binding curves of NS1 proteins from TBEV-Hypr (green) and TBEV-Vs (gray) to immobilized human JAM-A, measured by ELISA across three independent experiments. NS1 at various starting concentrations (**[A]** = 1 µg/mL, **[B]** = 1 µg/mL or **[C]** = 1µg/mL) was titrated in 2-fold serial dilutions. KD values are indicated. Data points represent mean OD450 ± SD from duplicate wells. Curves were fitted using a one-site binding model. **(D)** Quantification of NS1 binding at 0.01 µg/mL and 0.25 µg/mL to JAM-A. Bars represent mean OD450 ± SD. **(E–G)** Binding curves of Hypr and Vs NS1 to immobilized human ZO-1 across three independent ELISA experiments with variable NS1 starting concentrations (**[E]** = 0.5 µg/mL, **[F]**= 1 µg/mL or **[G]**= 1 µg/mL). NS1 was titrated in 2-fold serial dilutions. Data represent mean OD450 ± SD from duplicate wells. Fitted curves and KD values are shown. **(H)** OD450 values at 0.01 µg/mL and 0.25 µg/mL NS1 concentrations. Bars represent mean ± SD.

Binding assays with ZO-1 revealed minimal interactions for either NS1, consistent with the absence of predicted structural interfaces (Figure. 5E-G, Table 1). These results reinforce the biological irrelevance of direct NS1-ZO-1 binding, despite ZO-1 mislocalisation being observed in NS1-treated cells in prior studies ^21, 32^

Together, these data demonstrate that TBEV NS1 engages JAM-A in a strain-specific manner, with only the virulent Hypr variant showing direct binding. These findings support a model in which direct NS1-JAM-A interaction may contribute to blood–brain barrier disruption and neuroinvasion in virulent TBEV strains (Graphical illustration).

## Discussion

Despite the high conservation of orthoflaviviruses NS1 across both tick-borne and mosquito-borne members, emerging evidence suggests divergent functional roles in pathogenesis shaped by vector lineage, host tropism, and strain-specific viral factors ^4, 6–9, 32–34^. This study provides mechanistic insight into NS1 interactions that could relate to strain-specific neuropathogenicity in TBEV, dependent on NS1 interactions with BBB tight junction proteins. A key finding is the direct, high-affinity interaction between NS1 from the neurovirulent Hypr strain and the adhesion molecule JAM-A, which was not the case for the low-virulence Vasilchenko strain. This differential binding was demonstrated by biochemical validation.

Given JAM-A’s established role in maintaining BBB integrity and regulating immune cell transmigration ^35, 36^, its engagement by Hypr NS1 likely contributes to paracellular permeability and endothelial destabilisation. This points to a targeted mechanism of BBB disruption distinct from the generalised vascular leakage seen in mosquito-borne DENV and ZIKV, where NS1 induces endothelial dysfunction via pattern recognition receptor activation (e.g., TLR4) or complement pathways ^3, 33^.

Unlike DENV NS1, which interacts with ZO-1 and claudins to broadly destabilize endothelial junctions ^21^, TBEV NS1 appears to selectively engage claudins, with Hypr NS1 showing additional binding to JAM-B. This suggests that BBB disruption in tick-borne orthoflaviviruses infection may rely more on precise modulation of endothelial interfaces critical for CNS entry, rather than pan-junctional disassembly. Our data support the hypothesis that Hypr NS1-JAM-A interaction facilitates early neuroinvasion, potentially serving as a gateway rather than a consequence of systemic vascular damage.

Structurally, NS1 is a conserved dimer/trimer comprising β-roll, wing, and β-ladder domains ^1, 2^. Comparative analyses reveal that the 22 amino acid differences between Hypr and Vs NS1 localize predominantly to the wing and β-ladder domains–regions implicated in host membrane interactions and immune modulation. In other orthoflaviviruses, these domains mediate lipid binding, immune complex formation, and cell surface anchoring ^37, 38^. The clustering of strain-specific substitutions in these surface-exposed regions supports a model in which minor sequence variations reshape host interaction profiles without altering NS1’s overall architecture.

This concept aligns with broader patterns in orthoflaviviruses evolution. For example, a few substitutions in NS1 from ZIKV (Asian vs. African lineages) or WNV (Lineage 1 vs. 2) are sufficient to alter neurotropism, likely via changes in host protein or membrane interactions ^28, 39, 40^. Our findings suggest similar adaptive pressures in tick-borne orthoflaviviruses, where subtle NS1 modifications may fine-tune endothelial engagement and influence neuroinvasive potential.

The absence of JAM-A binding by Vs NS1 underscores the existence of alternative CNS entry mechanisms. Although Vs is less neurovirulent in murine models, human infections can result in persistent neurological symptoms and chronic inflammation ^41^. This suggests that Vs pathogenesis may involve gradual neuroinvasion via immune cell trafficking, receptor-mediated transcytosis, or inflammation-induced paracellular leak, mechanisms that do not require direct NS1-JAM-A interaction.

Interestingly, Vs NS1 exhibited modest binding to Claudin-5 and Hypr NS1 to Claudin-10, key components of BBB tight junctions ^31, 35^. However, the physiological relevance of these interaction remains uncertain. Claudin extracellular loops are short and structurally constrained, and direct NS1–claudin interactions have not been conclusively demonstrated *in vivo*. It is more plausible that claudin disassembly during TBEV infection occurs secondarily via inflammatory signalling, as proposed in DENV models involving TLR4 activation and cytokine release ^42^. This is consistent with findings in other neurotropic viruses, such as JEV, where cytokine-mediated modulation of claudins -not direct viral protein binding - drives BBB permeability changes ^43^.

In summary, this study provides mechanistic evidence that NS1-mediated engagement of endothelial junction proteins, particularly JAM-A, underlies strain-specific neuroinvasion in TBEV. By integrating computational structure prediction, evolutionary sequence analysis, and *in vitro* validation, we offer a grounded and testable model for differential pathogenesis. Future studies should investigate the *in vivo* mechanisms of NS1-mediated junctional disruption and assess how naturally occurring mutations influence these interactions in emerging strains. Given the expanding geographic range of both tick- and mosquito-borne orthoflaviviruses under climate change ^44–46^, monitoring NS1 sequence evolution may provide early indicators of shifts in virulence or transmission potential, informing diagnostic development and vaccine design.

## Acknowledgements

We thank the Wellcome Trust CDA Fellowship Scheme (226484/Z/22/Z). The funder had no role in the conceptualization, design, collection of data/analysis, nor the decision to publish the manuscript. We also thank Sam Driver (University of Leeds, UK) for technical assistance.

## Contributions

J.P., J.C., J.L., S.K., J.H.T., N.G, conducted the study or collected data. J.H.T., I.M.J., N.G. analysed or interpreted the data. N.G. conceived, supervised or designed the study, and drafted the manuscript. J.H.T., I.M.J., N.G. critically revised the article. All authors provided final approval of the version to be published.

